# An Accessible 3D HepG2/C3A Liver Spheroid Model Supporting the Complete Intrahepatocytic Lifecycle of *Plasmodium falciparum*

**DOI:** 10.1101/2024.11.06.621813

**Authors:** Claire H Caygill, Salwa Omar Alqurashi, Adriana Adolfi, Jessica Carson, Angelika Sturm, Daniel S Evans, Jess B Jinks, Koen J Dechering, Lisa Reimer, Shaun H Pennington, Parveen Sharma, Stephen A Ward, Giancarlo A. Biagini

**Affiliations:** Liverpool School of Tropical Medicine, UK; TropIQ Health Sciences, The Netherlands; University of Liverpool, UK

**Keywords:** Malaria, *Plasmodium*, HepG2/C3A, Spheroids

## Abstract

Current liver-stage *Plasmodium falciparum* models are complex, expensive, and largely inaccessible, hindering research progress. Here, we show that a 3D liver spheroid model grown from immortalized HepG2/C3A cells supports the complete intrahepatocytic lifecycle of *P. falciparum*. Our results demonstrate sporozoite infection, development of exoerythrocytic forms, and breakthrough infection into erythrocytes. The 3D-grown spheroid hepatocytes are structurally and functionally polarised, displaying enhanced albumin and urea production and increased expression of key metabolic enzymes, mimicking in vivo conditions – relative to 2D cultures. This accessible, reproducible model lowers barriers to malaria research, promoting advancements in fundamental biology and translational research.

## Introduction

Malaria continues to be a global health emergency. Despite intensive efforts, the WHO’s 2023 World malaria report estimated that the number of global malaria cases in 2022 remains stubbornly high at 249 million, exceeding pre-COVID-19 pandemic levels in 2019 (World malaria report 2023).

Chemotherapy continues to be a cornerstone to the multifaceted approach for malaria control and elimination and whilst the past two decades has delivered a robust pipeline of promising candidate drugs and several new treatments that act on the asexual blood stages (categorised as target candidate profile 1 [TCP-1]), there has been little progress in the development of effective therapies active against liver-stage hypnozoites (*Plasmodium vivax* and *P. ovale*) or hepatic *P. falciparum* schizonts (TCP-3 and TCP-4, respectively)(Siqueira-Neto *et al*., 2023).

The establishment of a stable *in vitro* culture system for the asexual blood stages of *P. falciparum* (Trager and Jensen, 1976) has been vital in unlocking our understanding of the intraerythrocytic stages of malaria infection and in advancing antimalarial drug discovery efforts for malaria treatment (TCP-1). Conversely, the relative paucity in our understanding of the biology of the hepatic stage of development and limited TCP3/4 pipeline has, in part, been attributed to the difficulty of establishing *in vitro* human liver stage malaria models, with current methods deemed expensive, laborious and impractical (Siqueira-Neto *et al*., 2023).

Initial studies attempting to establish stable *in vitro* models of malaria life stages using immortalized hepatoma cell lines, such as HepG2, HSS-102, and HC-04, achieved limited success and were unable to demonstrate breakthrough infection into erythrocytes. While these human hepatoma cell lines, allowed partial development of *Plasmodium* parasites, significant limitations persisted. For instance, HepG2-A16 cells could support sporozoite invasion but failed to sustain full parasite development. Similarly, HC-04 cells, though capable of supporting the liver stages of *P. falciparum* and *P. vivax*, exhibited reduced parasite viability and did not facilitate merozoite egress into erythrocytes due to their limited culture lifespan (Chattopadhyay *et al*., 2010; Tweedell *et al*., 2019; Valenciano *et al*., 2022).

More sophisticated approaches have achieved greater success in developing *in vitro* models for malaria research. Techniques such as using induced pluripotent stem cell (iPSC)-derived hepatocyte-like cells (iHCLs), immortalised hepatocyte-like cell (imHC) lines, co-culture systems, 2D primary human hepatocyte (PHH) monolayer systems, and 3D cell cultures have shown considerable promise (Valenciano *et al*., 2022). These advanced models have allowed large-scale drug screening campaigns (Roth *et al*., 2018; Maher *et al*., 2021), and have improved our understanding of *Plasmodium* liver stage biology. However, challenges remain, including limited expression of drug metabolism enzymes, difficulty in maintaining cultures long-term, and low sporozoite infection rates. Additionally, these advanced technologies are not widely accessible, particularly to laboratories in malaria-endemic low- and middle-income countries (LMICs) that often face resource and capability constraints. This disparity highlights the need for more accessible and robust *in vitro* models to support global malaria research efforts.

Multiple studies have shown that immortalised hepatocarcinoma cell lines cultured in 3D have superior liver-specific functionality when compared to monolayer cultures (Cox *et al*., 2020). A simple, high-throughput method of generating 3D cultures is through the creation of ‘spheroids’. Unlike more complicated 3D cell culture techniques, spheroids are simple to generate, in-expensive to culture, reproducible and may be analysed using a variety of methods (Cox *et al*., 2020).

HepG2/C3A cells are a derivative of the HepG2 cell line and have been shown to display greater contact inhibition and higher levels of liver-specific proteins and metabolic enzymes (Štampar *et al*., 2020, Kelly, JH US Patent 5290684, 1990). Here, we investigated whether a previously reported method of generating 3D liver spheroids from the immortalised HepG2/C3A cell line which demonstrated *in vivo*-like morphology and phenotype (Gaskell *et al*., 2016; Štampar *et al*., 2020), could be used to support the complete intrahepatocytic lifecycle of *Plasmodium*. Data is presented that demonstrate the HepG2/C3A liver spheroid model supports initial *P. falciparum* sporozoite infection, the development of exoerythrocytic forms (EEFs), and the subsequent egress of merozoites and breakthrough infection of erythrocytes. These data are discussed in the context of a method that is reproducible, inexpensive and accessible.

## Materials and methods

### HepG2/C3A 2D Cell Culture

HepG2/C3A cells were maintained in EMEM (LGC standards, Middlesex, UK), supplemented with 10% FBS (Sigma, UK) and 1% penicillin/streptomycin (Fisher Scientific, UK) at 37°C with 5% CO^2^. Cell culture medium was replaced every 3 to 4 days and cells were not allowed to exceed 80% confluency. Upon reaching 80% confluence, HepG2/C3A cells were washed once with PBS and incubated 0.25% (w/v) Trypsin EDTA (Fisher Scientific, UK) for 5 minutes at 37°C to detach the cells from the flask. After 5 minutes, cell culture media to was added and centrifuged at 173 x g for 5 minutes. Supernatant was then removed, remaining pellet resuspended in cell culture media and cell number counted with hemacytometer. HepG2/C3A cells were then seeded into fresh flasks at a seeding density of 2×10^5^ cells per mL. For albumin and urea comparative studies to 3D spheroids, HepG2/C3A cells were seeded 1000 cells per well (matching spheroid seeding density, described below) in 96-well cell culture plates in cell culture media and incubated at 37°C with 5% CO^2^ for 20 days.

### HepG2/C3A spheroid formation and growth

HepG2/C3A spheroids were grown using the liquid overlay technique, described elsewhere (Gaskell *et al*., 2016; Delves *et al*., 2016). In brief, 100μl of sterile 1.5% agarose in EMEM was added per well to flat bottom 96-well cell culture plates. Plates were then allowed to cool and solidify for 3 hours before and stored at 4°C upside down to prevent condensation for up to one month. Agarose plates were pre-warmed to 37°C and HepG2/C3A cells were seeded at 1000 cells per well in 100μl EMEM. Seeded plates were then centrifuged at 173 x g for 5 minutes and incubated for 72 hours at 37°C with 5% CO_2_ to allow formation of spheroids. Media changes were conducted every 2-3 days. Spheroid growth was monitored for 21 days and representative images taken using Echo Revolve R4 microscope with 20× objective. Images were analysed using ImageJ software to obtain spheroid diameter.

### Quantification of protein, albumin, and urea production

Cell lysates were obtained by disrupting 2D monolayer cultures and HepG2/C3A spheroid cultures on day 5, 9, 12, 14 and 21 using lysis buffer (250mM sucrose, 50mM Tris-HCl pH7.4, 5mM MgCl2, 1mM β-mercaptoethanol and 1% Triton X-100) and stored at −80°C. Protein concentrations from spheroid and monolayer cell lysates were quantified using Bio-Rad Protein Assay (Bio-Rad Laboratories), according to the manufacturer’s instructions.

Supernatants for spheroid and 2D cultures were also collected on days 4, 8, 12, 16 and 20 and stored at −80°C until required for quantification of albumin and urea. Albumin and urea were quantified using the Albumin Human ELISA Kit (Abcam) and the Urea Assay Kit (Abcam), respectively, according to manufacturer’s instructions. Albumin and urea data were normalised to total 2D or 3D spheroid protein content, as measured by Bio-Rad Protein Assay, described above.

### Measurement of CYP3A4 and CYP2D6 activity

CYP3A4 and CYP2D6 activity was measured using P450-GLO assay (Promega) on day 4 of 2D and 3D HepG2/C3A cultures, according to manufacturer’s instructions. Data was normalised based on the viable cell mass of day 4 2D and 3D spheroid cultures to account for any differences between cultures. This was measured by CellTitre-Glo Luminescent Cell Viability Assay (Promega), according to the manufacturer’s instructions. In brief, CellTitre-Glo reagent was added to 2D and 3D cultures, followed by mixing on an orbital shaker for 2 minutes to induce cell lysis and then incubated for 15 minutes at room temperature. Luminescence was then measured using POLARstar OMEGA plate reader (BMG Labtech).

### Immunofluorescence analysis of spheroids

Spheroids were fixed with 4% paraformaldehyde for 1h at 4°C, washed three times with PBS and permeabilised with 0.5% Triton X-100 in Tris-Buffered Saline with 0.05% Tween20 (TBST) overnight at 4°C. Spheroids were then blocked with 0.1% Triton X-100/5% BSA in TBST for 2h at room temperature. Primary antibodies monoclonal antibody 7.2 anti-GAPDH (1:100, obtained from The European Malaria Reagent Repository [http://www.malariaresearch.eu]), rabbit anti-HSP70 (*P. falciparum*) polyclonal (1:200, StressMarq Biosciences inc., SPC-186D) or rabbit recombinant monoclonal multidrug resistance-associated protein 2 (MRP2) antibody (1:200, Abcam, ab172630) were diluted in 0.1% Triton X-100/1% BSA in TBST and incubated with spheroids overnight at 4°C. Spheroids were washed three times with 0.1% Triton X-100 in TBST and incubated with Goat anti-Rabbit IgG (H+L) Cross-Adsorbed Secondary Antibody, Alexa Fluor™ 568 (1:1000, ThermoFisher Scientific, A-11011), Goat anti-Mouse IgG (H+L) Cross-Adsorbed Secondary Antibody, Alexa Fluor™ 488 (1:1000, ThermoFisher Scientific, A-11001), goat anti-Rabbit IgG (H+L) Cross-Adsorbed Secondary Antibody, Alexa Fluor 647 (1:1000, Thermo Fisher Scientific, A-21244) or Alexa Fluor 488 mouse monoclonal Anti-PGP9.5 (P-Glycoprotein 9.5 [Pg-P]) antibody (1:1000, Abcam, ab197733) in in 0.1% Triton X-100/5% BSA in TBST and incubated overnight at 4°C. DAPI, to detect cell nuclei (1:5000, ThermoFisher Scientific, 62248), and/or phalloidin for detection of F-actin filaments (1:250, ThermoFisher Scientific, A12380) in 0.1% Triton X-100/5% BSA in TBST was added to the spheroids and incubated for 1h at room temperature in the absence of light. Finally, spheroids were mounted with Prolong Gold (Life Technologies, P36930) onto a glass microscope slide with round, 13mm coverslips (Agar Scientific Ltd, UK). Images were acquired at 40× objective using a Zeiss LSM880 confocal microscope and analysed with Zeiss Zen Blue software (Zeiss, Germany).

### Statistical analyses

Multiple unpaired t-tests with Holm-Šídák correction for multiple comparisons and single unpaired t-tests were performed using GraphPad Prism version 5.01 (GraphPad Software, San Diego, USA).

### Infection of HepG2/C3A spheroids with P. falciparum NF54

*P. falciparum* NF54 parasites were maintained in continuous culture using established method (Trager and Jensen, 1976). Sexual stages of the parasites were cultured at 4% haematocrit in complete medium (RPMI 1640 with L-glutamine, 25mM HEPES pH7.4 (Gibco), 2g/L sodium bicarbonate, 50mg/ml hypoxanthine (Sigma) and 10% human heat-inactivated AB+ serum) from 0.5% mixed stage asexual cultures. Media was changed daily for 16-17 days and appearance of gametocyte stages was monitored every 48-72 h by microscopic analysis of thin blood smears (Delves et al., 2016).

Female *Anopheles coluzzii* mosquitoes aged between 5-7 days were deprived of sugar for 18-20h prior to infection. The standard membrane feeding assay was used to provide the mosquitoes with *P. falciparum* NF54 cultures containing stage V gametocytes for 30min (Tripathi et al., 2020). Afterwards, engorged mosquitoes were incubated in insectary conditions and offered ad libitum 10% sucrose solution. On day 16 post-infection with *P. falciparum* NF54, mosquitoes were collected and washed with 70% v/v ethanol and dissected to obtain salivary glands. Salivary glands were crushed in PBS and sporozoites stored in liquid nitrogen until required.

HepG2/C3A spheroids, grown as described above, were infected with *P. falciparum* NF54 20×10^4^ sporozoites per spheroid and incubated at 37°C with 5% CO_2_. On day 4 post-infection HepG2/C3A spheroids were fixed with 4% paraformaldehyde for 1h at 4°C and liver stage parasites detected by immunofluorescent staining of *P. falciparum* GAPDH and HSP70, as described above. The percentage of infected hepatocytes was calculated semi-quantitatively by counting the number of GAPDH or HSP70 positively fluorescing cells and dividing by the total cell number, as visible from DAPI staining of representative images from 4 spheroids, from 2 experimental replicates. Media changes were performed every 48 – 72 hours to maintain HepG2/C3A spheroids.

To access blood-stage breakthrough, human O+ erythrocytes were introduced to *P. falciparum* NF54 infected HepG2/C3A spheroids at a final haematocrit of 0.1% in complete EMEM and incubated at 37°C, with 5% CO_2_. On day 8 and day 12, 50 μL samples were taken from each well and gassed with 3% O_2_, 4% CO_2_ and 93% N_2_ and replenished with EMEM and human erythrocytes. To monitor asexual stage development, thin blood smears were prepared from the blood samples and stained with 10% Giemsa. Slides were examined under oil immersion at 100× using a Zeiss LSM880 confocal microscope (Zeiss, Germany). A schematic of the timeline for infection and sample collection is shown in Figure 1.

**Figure 1.**
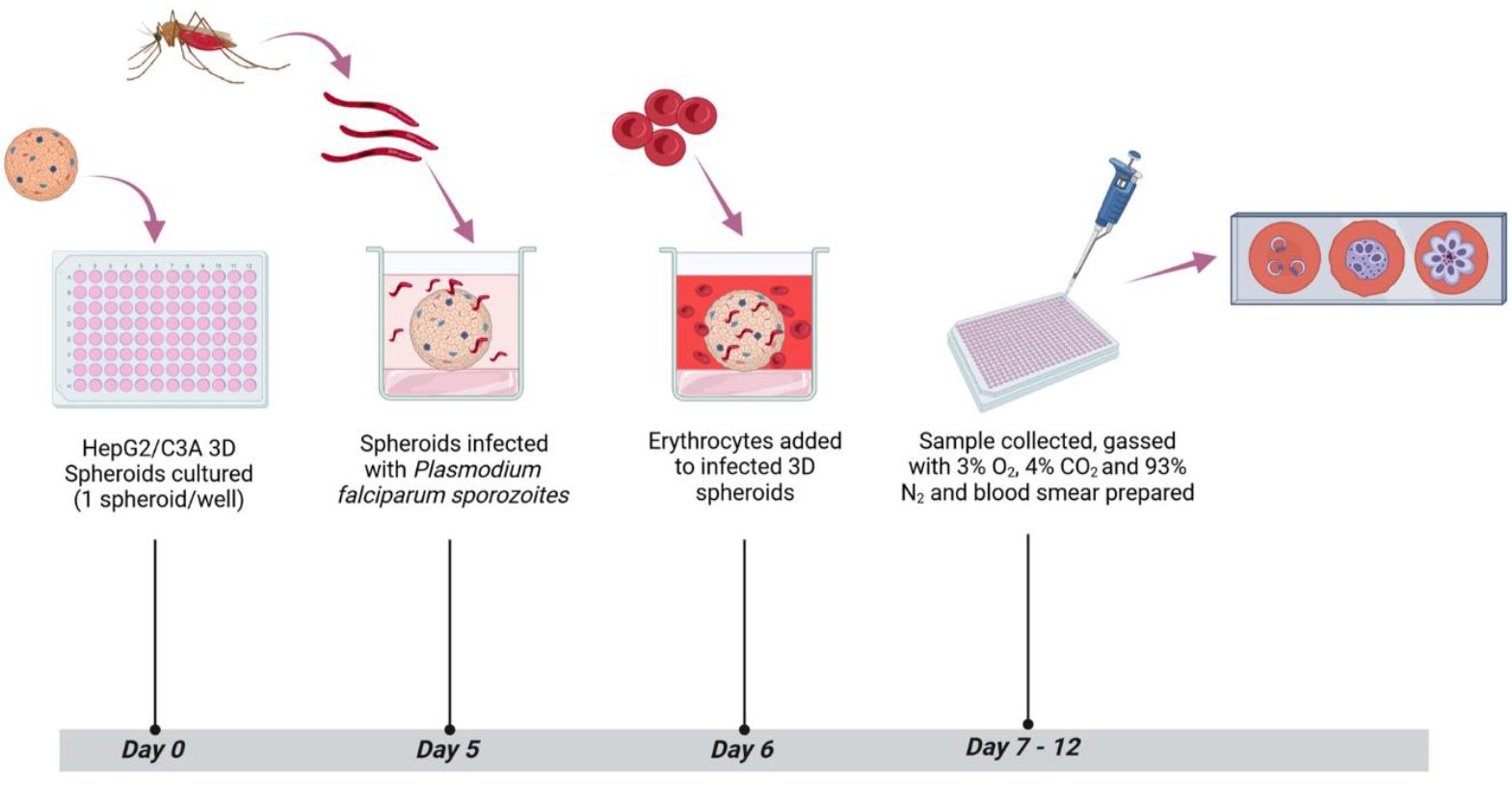
Schematic of HepG2/C3A 3D spheroid infection with *P. falciparum*, sample collection and assessment of blood-stage breakthrough timeline.

## Results

### HepG2/C3A spheroid formation and spheroid structure

To confirm the reproducibility of HepG2/C3A spheroid generation, assays to characterise spheroid formation and liver-like functions were undertaken, prior to infection with *Plasmodium*, as reported previously (Gaskell *et al*., 2016; Štampar *et al*., 2020). Spheroids aggregated by day 5 (Figure 2A) and their diameter increased between day 5 and day 21 from 210 μM ± 6.07 to 483.16 μM ± 19.26, respectively (Figure 2B). F-actin filaments and hepatocyte transporters (Pg-P and MRP2) were visualised by immunofluorescence staining on days 4 and 15. HepG2/C3A spheroid polarisation was confirmed by colocalization of F-actin with Pg-P and MRP2 (Figure 3). 2D monolayer HepG2/C3A cells were also imaged on day 4 and 15 and although F-actin filaments were visible at both time points, MRP2 and Pg-P transporters were not observed (Figure 3).

**Figure 2.**
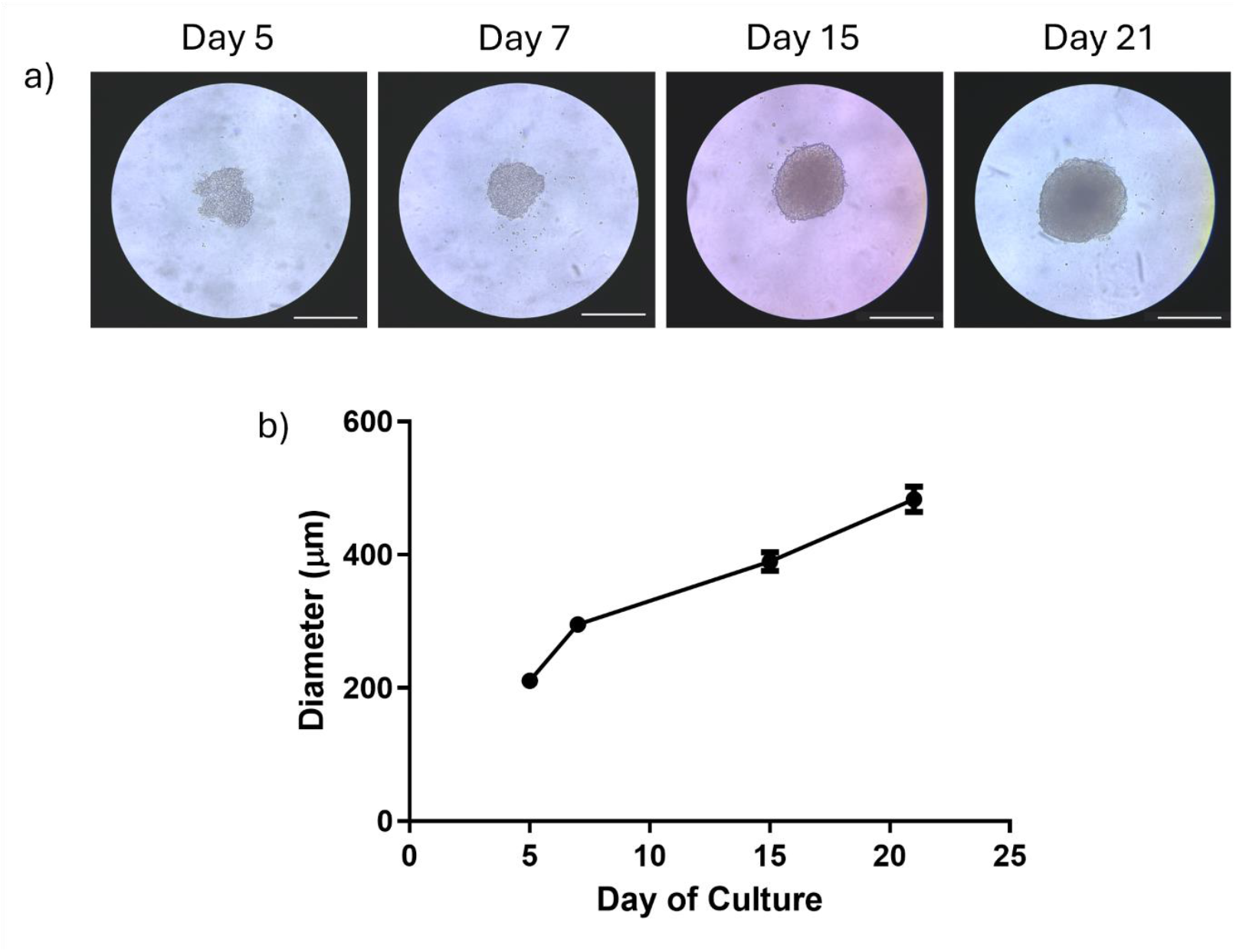
HepG2/C3A spheroid growth. HepG2/C3A cells were seeded at 1000 cells per well and incubated for 21 days to allow spheroid formation. Growth was observed and images acquired using Echo Revolve R4 microscope with 20× objective on days 5, 7, 15 and 21. Representative images are displayed, white scale bar represents 380μm (a). Images were analysed using ImageJ software to obtain spheroid diameter (b). Data represent the mean ± SEM of n=3 biological replicates, conducted in triplicate.

**Figure 3.**
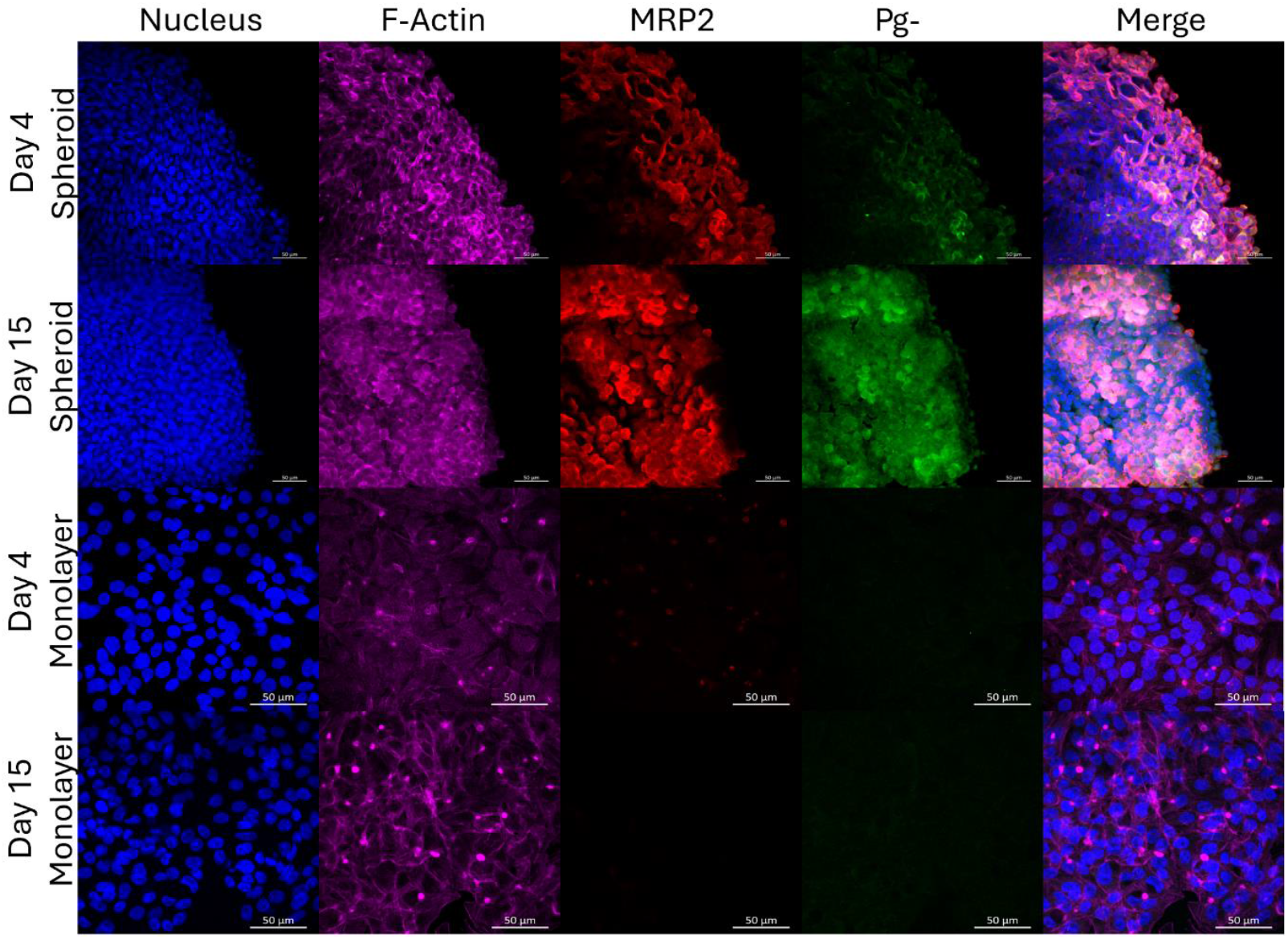
Hepatocyte transporters Pg-P and MRP2 expressed in HepG2/C3A 3D spheroids, but not in 2D culture. Secondary structures were visualised by immunofluorescence staining of HepG2/C3A 2D monolayer and 3D spheroid cultures on day 4 and 15. Phalloidin stain was used to visualise F-actin (purple) and canalicular transporters MRP2 (red) and Pg-P (green) were visualised with rabbit recombinant monoclonal MRP2 antibody (1:200, Abcam, ab172630), with goat anti-Rabbit IgG (H+L) Cross-Adsorbed Secondary Antibody, Alexa Fluor 647 (1:1000, Thermo Fisher Scientific, A-21244) and Alexa Fluor 488 mouse monoclonal Anti-PGP9.5 (Pg-P) antibody (1:1000, Abcam, ab197733), respectively. HepG2/C3A nuclei were stained with DAPI (blue). Representative images are shown of n=3 biological replicates, images were taken using Zeiss LSM880 confocal microscope, scale bar represents 50μm.

### Confirmation of liver-like function in spheroids

Spheroid albumin and urea production was quantified over 20 days and compared with 2D monolayers (Figure 4A and 4B). Albumin production in 2D monolayer cells remained low throughout the 20-day time course, with maximum albumin production observed on day 12 (1343.77 ng/mg ± 290.77 albumin). After day 12, albumin production in monolayer cells further reduced to 360.60 ng/mg ± 23.12 by day 20. 3D spheroids, however, demonstrated increases in albumin levels throughout the 20-day period, with the greatest albumin production observed on day 20 (5303.63 ng/mg ± 329.63). Significantly more albumin was produced in 3D spheroids, compared to 2D monolayer cells from day 8 onwards (Figure 4A). Similarly to albumin, urea production in 2D monolayer cells peaked on day 12 (2.58 nmol/mg ± 0.27) and decreased afterwards to 0.784 nmol/mg ± 0.029 by day 20. In contrast, 3D spheroid urea production increased over 20 days from 10.49 nmol/mg ± 0.77 on day 4 to 19.77 nmol/mg ± 0.522 on day 20. Significantly more urea was produced from 3D spheroids compared to 2D monolayer cells at every time point (Figure 4B).

**Figure 4.**
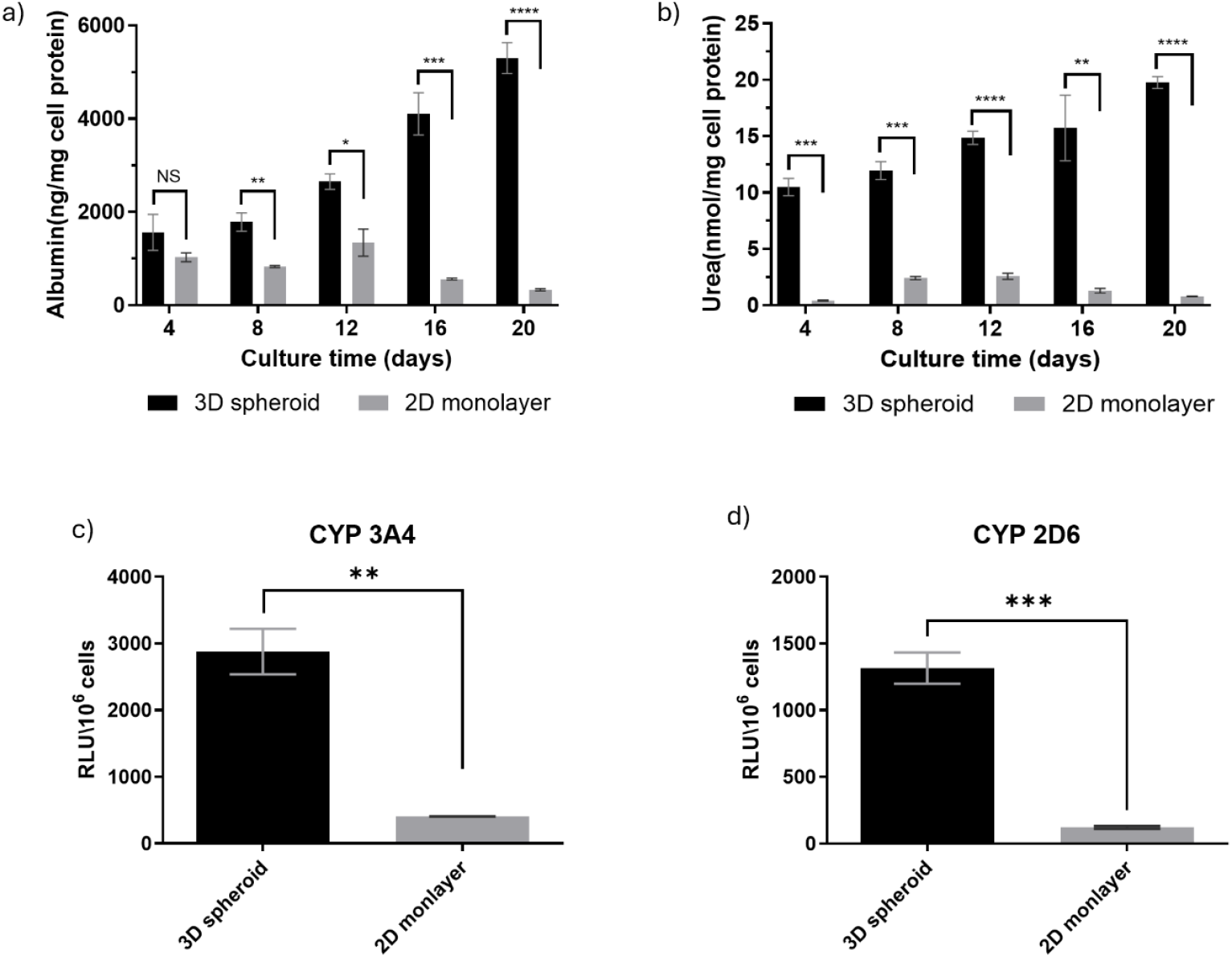
Functional characterisation of HepG2 3D spheroids and 2D cultures. Quantification of a) Albumin and b) Urea from HepG2/C3A grown in 2D and as 3D spheroids supernatants, normalised by mg cell protein on days 4, 8, 12, 16 and 20. Data were analysed by multiple unpaired t-tests with Holm-Šídák correction for multiple comparisons. Comparison of c) CYP 3A4 and d) CYP 2D6 expression from HepG2/C3A grown in 2D and as 3D spheroids on day 4, normalised by cell number, expressed as relative light units (RLU) per 10^6^ cells. Data were analysed by unpaired t-test. All Data represents the mean ± SEM of three independent experiments, performed with two technical replicates. * = p<0.05, ** = p<0.01, *** = p<0.001.

To access liver metabolism in HepG2/C3A 3D and 2D cultures, CYP3A4 and CYP2D6 activity was measured. Both enzymes have significant roles in drug metabolism (Tornio and Backman, 2018; Zhao *et al*., 2021) and would therefore be essential for any future works characterising anti-malaria drug efficacy. CYP3A4 activity was 7-fold higher and CYP2D6 10-fold greater in 3D spheroids (Figure 4C; 2878 ± 2.6 RLU/106 cells, 1315 ± 2.0 RLU/106 cells, respectively) compared to 2D culture (Figure 4D; 405 ± 0.7 RLU/106 cells, 122 ± 1.03 RLU/106 cell respectively).

### HepG2/C3A spheroid infection with P. falciparum NF54 sporozoites and Plasmodium asexual blood-stage breakthrough from HepG2/C3A spheroids to human erythrocytes

Spheroids were grown for 5 days and infected with 20×10^4^ *P. falciparum* NF54 sporozoites and incubated for a further 4 days, post-infection. Spheroids were then fixed and stained with antibodies to the sporozoite liver stage proteins GAPDH and HSP70 to confirm the presence of *P. falciparum* infection during the exoerythrocytic (hepatic) stages (Figure 5A and 5B). Uninfected control spheroids did not show staining of GAPDH or HSP70 (Figure 5C). Infection burden of liver spheroids was quantified in a semi-quantitative manner counting GAPDH or HSP70 positive cells as a percentage of infected hepatocytes relative to uninfected hepatocytes from representative spheroid images (Figure 5D). GAPDH and HSP70 indicated an infection burden of 28.7% ± 3.3% SEM and 12.3% ± 2.12% SEM, respectively.

**Figure 5.**
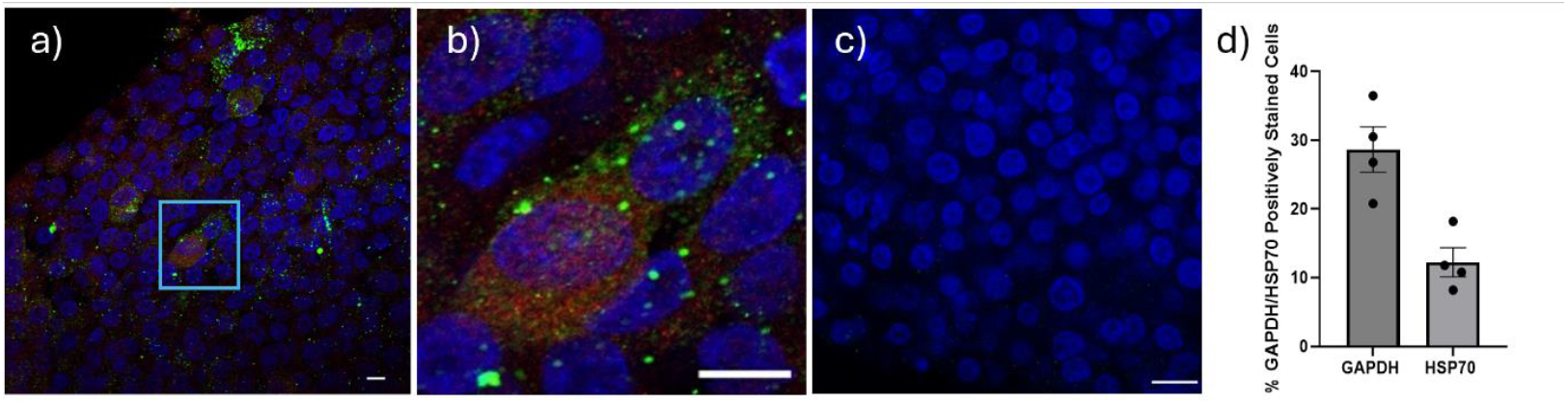
Assessment of HepG2/C3A 3D spheroid culture infection by *P*.*falciparum* NF54 sporozoites and asexual blood stage breakthrough from infected HepG2/C3A 3D spheroids. a) Liver stage parasites were detected by immunofluorescence on day 4 post-infection and stained with antibodies towards GAPDH (green [monoclonal antibody 7.2 anti-GAPDH 1:100, obtained from The European Malaria Reagent Repository [http://www.malariaresearch.eu]) and HSP70 (red [rabbit anti-HSP70 *P. falciparum* polyclonal, 1:200, StressMarq Biosciences inc., SPC-186D]), cell nuclei are visualised with DAPI stain (blue). Region of interest (ROI) is shown in light blue box. b) Close-up of ROI showing staining of GAPDH (green) and HSP70 (red) surrounding cell nuclei (blue). c) Uninfected HepG2/C3A spheroid control. d) Infection burden was expressed as the percentage of positively stained GAPDH or HSP70 hepatocytes relative to hepatocytes with no colocalised staining with DAPI. Data displayed is from n=2 biological replicates, carried out in duplicate. Error bars represent SEM. Images were obtained with Zeiss LSM880 confocal microscope (Zeiss, Germany) White scale bar represents 10 μm.

To access if HepG2/C3A infected spheroids were able to lead to blood-stage breakthrough, human erythrocytes were introduced on day 6, on two separate occasions and incubated till day 12. On day 8, a thin blood film was then prepared, and erythrocytes were observed to contain ring staged parasites (Figure 6A). After 12 days, another thin blood film was then prepared and trophozoite/early schizont parasites were seen (Figure 6B). Blood stage breakthrough was observed from all infected spheroids.

**Figure 6.**
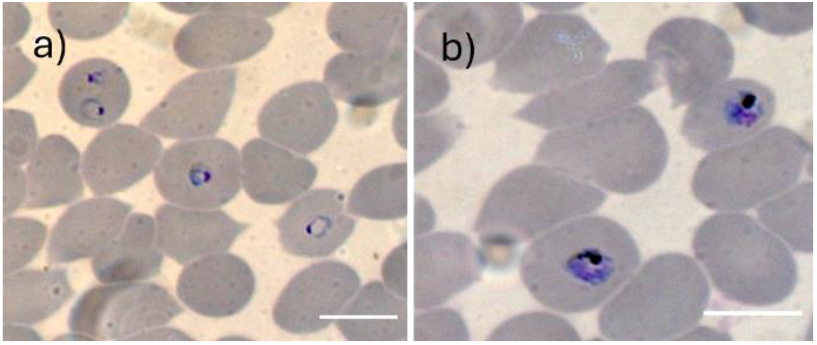
Asexual blood stage breakthrough from infected HepG2/C3A 3D spheroids. Human erythrocytes were introduced to infected HepG2/C3A cultures at a final haematocrit of 0.1% in complete EMEM and incubated for 6 days. Giemsa-stained thin blood films were performed on samples from day 8 and day 12 following infection with *P. falciparum* NF54 which were gassed with 3% O_2_, 4% CO_2_ and 93% N_2_, n=5 HepG2/C3A spheroids. On day 8 ring stage parasites were observed (a) and by day 12 trophozoite/early schizont parasites could also be detected (b). Slides were examined under oil immersion at 100× using a Zeiss LSM880 confocal microscope (Zeiss, Germany) White scale bar represents 10 μm

## Discussion

The HepG2/C3A liver spheroid model developed here represents a low-cost, accessible means to explore the complete intrahepatocytic lifecycle of *P. falciparum*, addressing significant limitations faced by many laboratories in adopting advanced *in vitro* models for malaria research.

3D cell culture technologies have increasingly been recognised for their ability to more accurately model *in vivo* cellular behaviour *in vitro* relative to 2D cultures. The hepatocytes within the 3D-grown HepG2/C3A spheroid were shown to be structurally and functionally polarised, recapitulating the *in vivo* condition (Godoy *et al*., 2013). Secondary structures composed of F-actin were observed forming enhanced networks throughout the spheroids, that were associated with Pg-P and MRP2, transporters localised to the canalicular membrane of hepatocytes, as described elsewhere (Gaskell *et al*., 2016). HepG2/C3A spheroids displayed enhanced synthesis and secretion of liver proteins, albumin and urea, as well as enhanced CYP metabolism, as exemplified by raised CYP 3A4 and CYP 2D6 expression, compared to monolayer cultures. CYP 3A4 and CYP 2D6 expression was then shown to increase over time in 3D spheroids. It should be noted that CYP metabolism comparison between monolayer and 3D spheroids was conducted on day 4 only, as accurate comparisons could not be assured as the spheroid size and density grew and full penetration of CYP luciferin pro-substrates could not be guaranteed. This could therefore have led to an underestimation of differential CYP expression, compared to 2D monolayers. These data support previous observations (Gaskell *et al*., 2016; Štampar *et al*., 2020) and confirm that cells cultured in 3D have superior functionality to those cultured in 2D (Cox *et al*., 2020; Tutty *et al*., 2022).

Previous attempts to create stable *in vitro* models of malaria life stages using immortalised hepatoma cell lines, such as HepG2, have had limited success and failed to demonstrate breakthrough infection into erythrocytes (Chattopadhyay *et al*., 2010; Tweedell *et al*., 2019; Valenciano *et al*., 2022). However, the structural and functional attributes of 3D-grown HepG2/C3A cells appear to offer the necessary physiological conditions for *P. falciparum* intrahepatocytic development. Our findings suggest these conditions are essential for completing the parasite’s lifecycle within hepatic cells.

The current study is limited as it is a qualitative assessment of blood-stage breakthrough from *P. falciparum* infected HepG2/C3A liver spheroids using a relatively large sporozoite infection burden. To fully benchmark this system against existing liver-stage models, a more extensive comparative study would be required in the future. The described HepG2/C3A 3D spheroids model also presents a unique opportunity to identify proteins involved in stage conversion in follow-on studies. These studies were beyond the scope of the presented article, however, one recent global gene expression study of human malaria parasite liver stages revealed significantly increased expression of genes in central metabolic pathways and redox homeostasis (Zanghi *et al*., 2023). It is noteworthy that previous studies have shown that HepG2/C3A 3D spheroids replicate the liver’s zonation due to oxygen and nutrient gradients (Gaskell *et al*., 2016; Cox *et al*., 2020). It will be important to determine whether the critical contributing factor to the permissiveness of the 3D spheroid model to intrahepatocytic parasite development is that the specific nutrient and redox needs of the parasite can be met by the unique spatiotemporal microenvironment of the spheroid.

In summary, the HepG2/C3A liver spheroid model lowers barriers to malaria research by offering a simpler and more accessible method to study the hepatic stages of *P. falciparum* infection. This model addresses a critical gap in malaria research tools and can significantly enhance our ability to study fundamental aspects of *Plasmodium* biology and develop effective liver-stage interventions. This approach not only broadens the scope of research possibilities but also provides a platform for high-throughput screening of antimalarial drugs and vaccines, promoting advancements in both fundamental biology and translational research.

## Supplementary material

N/A.

## Author’s contribution

CHC, SOA, AA, JC, AS, LR, DSE and JBJ conducted data gathering. CHC and SOA performed statistical analyses. SOA, AA, AS, LR, KJD, SHP, PS, SAW, and GAB contributed to the study design. CHC, SOA and GAB wrote the article. GAB conceived the study.

## Financial support

For the purpose of open access, the authors have applied a Creative Commons Attribution (CC BY) licence to any Author Accepted Manuscript version arising this submission. This study was supported by funding from the Medical Research Council (MR/W002248/1, MR/L000644/1 and MR/R015678/1) and by the Bureau Culture of Saudi Arabia under the directives of King Salman Bin Abdulaziz (SOA).

## Competing interests

The authors declare there are no conflicts of interest.

## Ethical standards

Not applicable

## Notes

### Competing Interest Statement

The authors have declared no competing interest.

### Summary of Updates

Figure 1 - new figure to show methodology Figure 2 - was supplementary, now in main document Figure 3 - additional images and control images added Figure 4 - 2D culture values added

